# Regulation of tensile stress in response to external forces coordinates epithelial cell shape transitions with organ growth and elongation

**DOI:** 10.1101/399303

**Authors:** Ramya Balaji, Vanessa Weichselberger, Anne-Kathrin Classen

## Abstract

The role of actomyosin contractility at epithelial adherens junctions has been extensively studied. However, little is known about how external forces are integrated to establish epithelial cell and organ shape *in vivo*. We use the *Drosophila* follicle epithelium to investigate how tension at adherens junctions is regulated to integrate external forces arising from changes in germline size and shape. We find that overall tension in the epithelium decreases despite pronounced growth of enclosed germline cells, suggesting that the epithelium relaxes to accommodate growth. However, we find local differences in adherens junction tension correlate with apposition to germline nurse cells or the oocyte. We demonstrate that medial Myosin II coupled to corrugating adherens junctions resists nurse cell-derived forces and thus maintains apical surface areas and cuboidal cell shapes. Furthermore, medial reinforcement of the apical surface ensures cuboidal-to-columnar cell shape transitions and imposes circumferential constraints on nurse cells guiding organ elongation. Our study provides insight into how tension within an adherens junction network integrates growth of a neighbouring tissue, mediates cell shape transitions and channels growth into organ elongation.

## Introduction

Epithelia give rise to the branching surface of lungs, convoluted villi of the gut and the protective layer of our skin. The diversity in epithelial function is matched by the diversity in epithelial cell shapes. Squamous, cuboidal and columnar cell shapes arrange in monolayers or stratify to support the function of the respective tissue. These shapes are determined by cell-intrinsic mechanical properties and force-generating mechanisms that the interplay between adhesion and cytoskeleton generates. However, during organogenesis or homeostatic maintenance, epithelial cells must also integrate forces which arise from growth or shape changes of neighboring tissues. These external forces can stretch, shear or compress tissues and alter cell shape. Thus, the balance between cell-intrinsic and external forces ultimately defines 3D cell shape [1]. The role of actomyosin contractility in bringing about cell-intrinsic shape changes, such as remodeling in the plane of adherens junctions, is well documented [2-6]. However, much less is known about how contractility is reinforced to resist external forces and thereby helps to prevent cell shape deformation. Similarly, little is known about how contractility is downregulated to mediate relaxation and cell shape transitions in response to external forces [7-9].

We wanted to specifically understand how external forces that stretch an epithelium are integrated by regulation of contractility at adherens junctions (AJ) and how the resulting tensile stress at AJs modulates transitions between cuboidal and columnar 3D cell configurations. We investigate this question in the *Drosophila* egg chamber consisting of a germline and a somatic follicle epithelium undergoing coordinated morphogenesis demarcated by 14 stages [10]. The germline consists of 15 ‘nurse’ cells and 1 oocyte, which grow in size until stage 11 when nurse cells rapidly transfer their cytoplasm to the oocyte during ‘nurse cell dumping’. The somatic follicle epithelium consists of ca. 850 initially cuboidal cells, which completely envelop the germline. Between stage 8 and 10A, about 50 cells specified by anterior fate patterning undergo a cuboidal-to-squamous shape transition to overlie the nurse cell cluster. Between stage 6 and 10A, about 800 posterior cells of main body and posterior terminal fate undergo a cuboidal-to-columnar shape transition to overlie the growing oocyte [11-13]. The differentiation of these diverse epithelial shapes contradicts the expectation that the epithelium subject to growth and expansion of the enclosed germline would accommodate the increase in shared surface area by uniform flattening. Although squamous cell flattening has been previously suggested to represent a compliant response to germline growth, cuboidal and columnar shapes may resist flattening by relatively higher apical stiffness [11]. However, little is known about how the follicle epithelium regulates its mechanical properties to integrate the growth of the enclosed germline with these cell shape transitions.

As follicle cells change shape in the absence of cell divisions or cell intercalation [10, 14], coordination of shape transitions within the epithelium can be exclusively attributed to cytoskeletal remodeling or turn-over of cell adhesion. Specifically, flattening of anterior cells is promoted by removal of Fasciculin 2 from lateral surfaces [15] and disassembly of E-cadherin (E-cad) dependent AJs [16]. Cuboidal-to-columnar shape transitions of posterior cells was thought to be driven by apical constriction [12, 17, 18]. However, an increase in lateral height driven by cellular growth fully accounts for columnarisation [11]. Several genetic studies indicate that Actin, non-muscle Myosin II (MyoII), Spectrins and Integrins are essential to maintain cuboidal and establish columnar cell shapes [19-23]. However, these studies have not revealed in detail how these components regulate epithelial behavior to integrate germline surface area expansion with cuboidal-to-columnar cell shape transitions.

Throughout all stages, the apical surface of the epithelium faces the interior germline (Fig S1A-A’’). Thus, any change in germline volume must be matched by a change in apical epithelial area. Apical MyoII has been proposed to withstand forces from germline growth prior to stage 6 and to promote cell divisions to compensate for germline surface growth during early stages [20]. However, how actomyosin is integrated with expansion of the apical surface after cell divisions ceased is not known. In contrast, the basal surface of the epithelium is facing the egg chamber exterior and deposits an ECM. Circumferentially polarized ECM fibrils have been proposed to cause egg chamber elongation during stage 2-8 by providing an external, patterned molecular corset channeling growth of the egg chamber in the anterior-posterior (A/P) axis [24-27]. This basal corset is thought to be supported by polarized Actin filaments [28] and oscillatory MyoII-driven contraction [29] to ensure organ elongation until stage 10A. However, the role of the apical surface in organ elongation has only recently received attention [30]. Here we provide new insight into regulation of AJs length and contractility to mediate epithelial relaxation during organ growth, to reinforce cuboidal and columnar cell shapes and to modulate organ shape.

## Results

### Main body and posterior terminal cell shape correlates with apposition to oocyte or nurse cell compartments

During stages 5 and 10A, the oocyte increases in size more rapidly than nurse cells (Fig 1A, E) [11]. As a result, an increasing number of follicle cells that are positioned over nurse cells, come into contact with the oocyte. Specifically, while only a few main body cells identified by *mirr* expression [13] contact the oocyte before stage 9, all main body cells contact the oocyte by stage 10A (Fig 1B,B’,F). Of note, all posterior terminal cells identified by *pnt* expression [13] contact the oocyte between stages 7 and 10A (Fig S1B).

**Fig 1:**
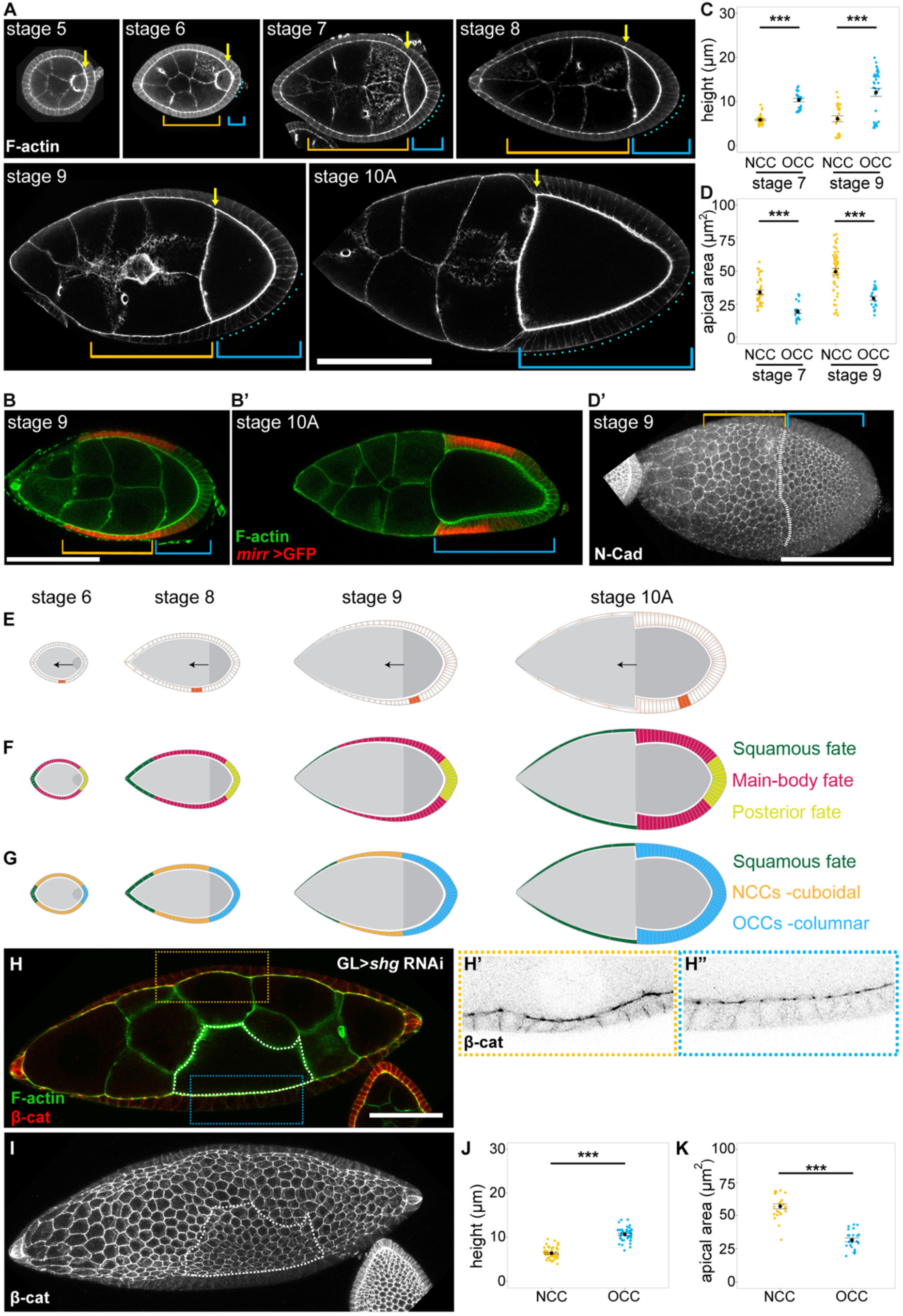
Main body and posterior terminal cell shape correlates with apposition to oocyte or nurse cell compartments. **(A)** Medial sections of stage 5 to 10A egg chambers stained for F-actin. Yellow arrows mark the position of the anterior oocyte boundary. Blue brackets and cyan dots identify columnar cells in contact with the oocyte called oocyte-contacting cells (OCCs, 4, 8, 10, 13 and 24 cells from stage 6 to 10A, respectively). Orange brackets identify cuboidal cells in contact with the nurse cells, which are not of squamous fate and will be called nurse-cell-contacting cells (NCCs). **(B-B’)** Medial sections of stage 9 and 10A egg chambers stained for F-actin (green) and expressing *UAS GFP* (red) driven by *mirr-Gal4* that is expressed in main-body-fated cells. Note that at stage 9, cells expressing *mirr-Gal4* are both cuboidal (NCCs, orange bracket) and columnar (OCCs, blue bracket), whereas by stage 10A all *mirr-Gal4* cells are columnar and OCCs. **(C-D’)** Quantifications of cell heights (C) and apical areas (D) of NCCs and OCCs at stages 7 and 9. Maximum projection of confocal sections of a stage 9 egg chamber to obtain *en face* view (see Fig S1A’) of AJs (D’). The egg chamber was stained for N-Cad to visualize apical NCC (orange bracket) and OCC (blue bracket) areas (D’). **(E-G)** Schemes of medial sections between stages 6 to 10A. 31 cells in a row span each half from the anterior to the posterior pole. Egg chambers in (E-G) can be superimposed according to stage. The position of the same three cells (orange, E) are tracked as the oocyte grows anteriorly (black arrows). The orange cells are initially in contact with nurse cells (stage 6 and 8) and as the oocyte moves anteriorly, they encounter the oocyte (stage 9 and 10A) (E). If future columnar cells (pink and yellow) are tracked by known developmental fate markers after stage 6 (F), neither main-body markers (pink, i.e. *mirr*) nor posterior terminal markers (yellow, i.e. *pnt*) track exclusively with cuboidal or columnar cell shape. Instead, cell shapes track with germline contact. Future columnar cells are cuboidal when over nurse cells (orange, NCCs), and columnar only upon contact with the oocyte (blue, OCCs) (G). **(H-K)** Medial sections to visualize cell height (H-H’’) and maximum projection for *en face* view of AJs (I) of a stage 9 egg chamber. The germline (GL) expresses RNAi targeting *E-cad (shotgun, shg)* resulting in a mispositioned oocyte. The oocyte was identified by the pronounced F-actin cortex and is framed by white dotted lines in (H) and (I). The egg chamber was stained for F-actin (green in H) and β-cat (red in H,H’-I). Orange (NCC) and blue (OCC) framed regions in (H) are shown at higher magnification in (H’,H’’), respectively. Note the differences in height (H’,H’’) and apical surface area (I) between NCCs and OCCs. Quantification of cell heights (J) and apical areas (K) of NCCs and OCCs in egg chambers with a misplaced oocyte. Graphs display mean±SEM. A WMW-test (C,D) and t-test (J,K) was performed. *** indicates p-value≤0.001. For sample sizes, see Table S2. Scale bar (A-D’)=100 μm, (H-I)=50 μm.

During stages 5 and 10A, all posterior terminal and main body cells transition from a cuboidal to a columnar aspect ratio [10, 11]. At each stage, cells in contact with the oocyte were taller in height and had a reduced apical surface area than follicle cells still contacting nurse cells (Fig 1C-D’). Thus, posterior terminal and main body cells in contact with the oocyte were columnar, whereas cells still contacting nurse cells were cuboidal. To account for this contact-correlated behavior of future columnar cells, we distinguish this population to either be nurse-cell-contacting cells (NCCs) or oocyte-contacting cells (OCCs) (Fig 1G). A previous study revealed that all future columnar cells grow equally in volume [11]. Thus, the contact-correlated differences in 3D cell shape represent a volume-conserved difference in the ratio of apical to lateral domains.

We wanted to understand if known oocyte-derived patterning signals, such as EGF-signaling, were sufficient to explain contact-dependent differences in NCC and OCC shapes. Expression of a constitutively active EGF receptor in all follicle cells prevents specification of anterior fates [13] and thus squamous cell flattening at stage 9 (Fig S1C). However, the height of main body follicle cells in contact with nurse cells was still smaller than of cells in contact with the oocyte (Fig S1C). Thus, while ectopic EGF signaling can alter anterior cell fate patterns, it is insufficient to directly promote columnar shape in main body cells still in contact with nurse cells (NCCs). Conversely, expression of a dominant-negative EGF receptor in main body follicle cells is sufficient to prevent formation of dorsal appendages (Fig S1D) [31] but did not prevent acquisition of columnar shape in main body cells in contact with the oocyte (OCCs) (Fig S1E,F). Thus, oocyte-derived EGF signaling is required for fate patterning but is neither sufficient nor necessary to promote the transition of main body follicle cells from cuboidal to columnar shape once they contact the oocyte.

To understand if contact to either oocyte or nurse cells was necessary for the differences in main body cell shape during stage 9, we analyzed conditions where the oocyte was mispositioned within the egg chamber [32, 33]. In these egg chambers, OCCs were still taller in height and had reduced apical areas than their nurse cell-contacting neighbors (Fig 1H-K). Thus, upon genetically separating acquisition of columnar shape from topological position within the egg chamber and from patterning by posterior pole cells, we find that cells lacking anterior fates only acquire columnar shape if in contact with the oocyte, while maintaining cuboidal shapes when still in contact with nurse cells.

### A medial shift of MyoII, emergence of junctional corrugations and remodeling of the Actin cortex reorganize the apical domain

As the follicle cell’s apical domain faces the germline, we speculated that nurse cell or oocyte signals may regulate the apical cortex and AJ network to control changes in apical follicle cell area and consequently, the shift in cellular aspect ratio from cuboidal to columnar shapes. We therefore analyzed localization of apical actomyosin and AJ markers between stages 6 and 9 to better understand how cell shape transitions may be regulated during these stages.

We found that MyoII-associated markers, such as the active phosphorylated form of the MyoII regulatory light chain (MRLC-1P) and the MyoII heavy chain (Zip (Zipper)) localized to AJs and the apical cortex at stage 6. Strikingly, by stage 9, all MyoII markers became depleted from AJs and medial MyoII enriched in the apical cortex (Fig 2A-D’’’’, Fig S2A,A’). Importantly, all follicle cells along the anterior-posterior (A/P) axis shifted MyoII to a medial localization by stage 9 (Fig S2B). Previous studies demonstrate cessation of apical-medial MyoII oscillations after stage 6 [30], an observation supported by a lack of apical-medial MyoII oscillations with a reported time period of 3 min in live imaging at stage 9 (Fig S2C). To understand how apical-medial MyoII may thus act, we visualized the subcellular organization of Actin. Total F-Actin staining and follicle-cell-specific expression of *utABD-GFP* revealed that apical-medial Actin filaments in the follicle cell cortex enriched between stages 6 and 9 (Fig 2E-F’’, Fig S2D). Of note, levels of apical Actin filaments were always higher in OCCs, likely reflecting the differentiation of apical microvilli (Fig S2E) [34]. Importantly, however, Actin unlike MyoII, was not excluded from junctions; in fact, Actin filaments radiated from cell-cell contacts formed by AJs. Strikingly, these pronounced changes to actomyosin organization at the apical cortex coincided with pronounced changes in AJ appearance. Between stages 6 and 9, follicle cell AJs started to exhibit gaps in E-cad/β-cat continuity and became more corrugated (Fig 2G-G’). We define corrugations as a larger than 1 ratio of the observed junctional length to that of a straight line between two cellular vertices. Specifically, the surplus junctional length connecting two cellular vertices compared to that of a straight line tripled between stages 6 and 9 (Fig 2G’’). Importantly, an increase in junctional corrugations was exclusively observed at the level of AJs; basolateral surfaces remained straight (Fig 2H-I’’). Both junctional depletion and medial shift of MyoII, as well as AJ corrugations, could also be observed during live imaging of egg chambers, excluding processing artifacts as a source for changes in apical-junctional architecture (Fig S2F-G’). Combined these observations reveal a pronounced reorganization of the apical-junctional cortex facing the growing germline surface during post-mitotic stages of egg chamber growth.

**Fig 2:**
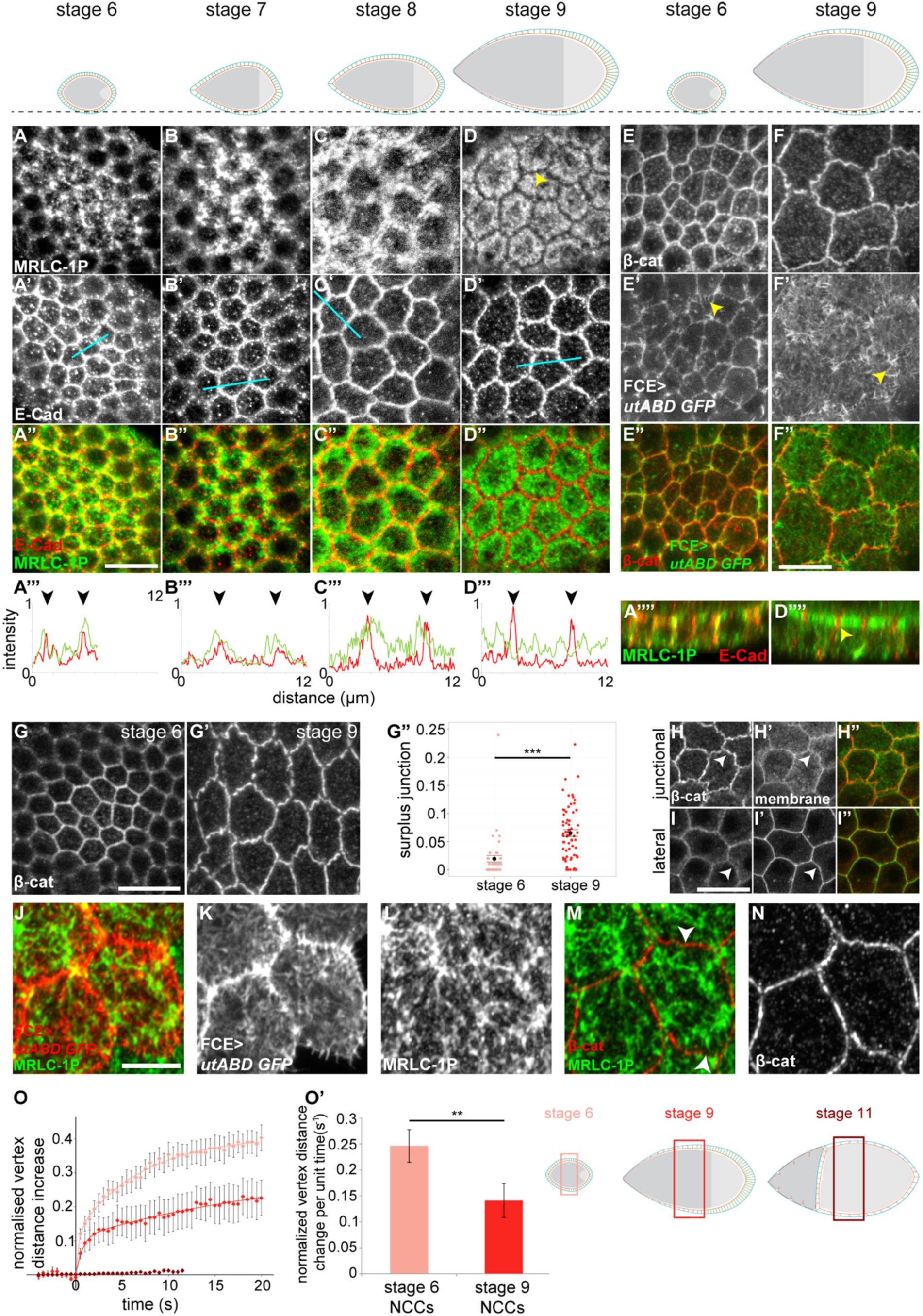
A medial shift of MyoII, emergence of junctional corrugations and remodeling of the Actin cortex reorganize the apical domain. **(A-D””)** *En face* AJ sections of egg chambers at stages 6 to 9 (A-D”) stained for MRLC-1P (A-D, green in A”-D”), E-cad (A’-D’, red in A”-D”). Black line across egg chamber schemes indicates position at which sections were acquired. Line profile plots (A’’’-D’’’) of MRLC-1P (green) and E-cad (red) fluorescence intensities along cyan lines in (A’-D’). Black arrowheads point to E-cad intensity peaks at AJs. XZ-reslices of confocal stacks from stage 6 and 9 epithelia (A’’’’,D’’’’) shown in A and D. Yellow arrowheads point to depletion of MRLC-1P from AJs. **(E-F”)** *En face* AJ sections of egg chambers at stages 6 to 9 stained for β-cat (E,F, red in E”,F”). Actin was visualized by follicle cell specific expression (FCE) of *UAS utABD-GFP* (E’,F’, green in E”,F”). Yellow arrowheads indicate Actin filaments positioned at an angle to junctions. **(G-G”)** *En face* AJ sections of NCCs at stage 6 (G) and 9 (G’) stained for β-cat. Note the increase in junctional corrugations. Quantification of surplus junctional length (G’’, see Exp. Proc.). Graphs display mean±SEM. n=46 (stage 6) and n=65 (stage 9) junctions at NCC positions in 3 egg chambers at each stage. WMW-test was performed. *** indicates p-value≤0.001. **(H-I”)** *En face* sections of NCC AJs (H-H”) and lateral interfaces (I-I”) at stage 9 stained for β-cat (H,I, red in H”,I”) and the plasma membrane marker PH-GFP (H’,I’, green in H”,I”). Arrowheads point to corrugations (H,H’) and their absence (I,I’). **(J-N)** *en face* super-resolution sections obtained by Airy-scan imaging of NCCs with FCE expression of *utABD-GFP* (K, red in J), stained for MRLC-1P (L, green in J,M) and β-cat (N, red in M). Arrowheads (M) point to MRLC-1P and actin filaments radiating from AJs and connecting to the medial actomyosin cortex. **(O-O’)** Graph (O) displays the normalized average increase in distance between vertices upon laser ablation of AJs (t=0) as a function of time in stage 6 (pink, n=10), stage 9 (red, n=10) and stage 11 (brown, n=8) egg chambers. Graph displays mean±SEM and a double exponential fit (see Exp. Proc.,Table S3,S4). Normalized initial vertex distance change per unit time (O’, see Exp. Proc.) of vertices at stage 6 NCCs and stage 9 NCCs. Graphs display mean±SEM. A two-sample t-test was performed. ** indicates p-value≤0.01. Egg chamber schemes illustrate position of follicle cells subject to laser cuts. Scale bars (A-I’)=10 μm, (J-N)=5 μm

To better understand how corrugating AJs may interface with apical-medial MyoII, we performed super-resolution microscopy of the apical cortex at stage 9. Medial Actin filaments visualized by follicle cell specific expression of *utABD-GFP* interdigitated with medial MyoII clusters (Fig 2J-L). Multiple examples of long medial MyoII filaments connecting to AJs could be observed (Fig 2L-N, Fig S2H-J), strongly indicating that the medial actomyosin cortex is connected to and acting on corrugating AJs. This rearrangement of medial-junctional architecture between stages 6 and 9 suggested profound alterations to contractile forces acting on AJs.

To specifically understand how tension at AJs may be affected by depletion of junctional MyoII, we measured recoil velocities upon laser ablation of AJs at stage 6 and stage 9 [35-37]. Strikingly, junctional tension decreased from stage 6 to stage 9 in main body follicle cells overlying nurse cells at either stage (Fig 2O,O’). This analysis revealed that, despite the dramatic growth and surface area expansion of the germline, which is expected to impose strain on the overlying epithelium, AJ tension decreases by stage 9. These observations suggest that the junctional depletion and medial shift of MyoII coincides with a reduction in AJ tension despite external forces imposed by germline growth. Strikingly, by stage 11, when oocyte-contacting cells need to accommodate the rapid increase of the oocyte surface during nurse cell dumping, junctional tension dropped to almost undetectable levels (Fig 2O). Combined, these results demonstrate that AJ tension within the closed sheet of the follicle epithelium is highly regulated. We speculate that junctional depletion of MyoII and increasing AJ length supports the developmentally coordinated reduction in AJ tension. Reducing AJ tension would promote relaxation in the plane of the junctional network and allow epithelial cells to adapt their apical-junctional area to the growing egg chamber surface.

### Levels of E-cad/β-cat, MyoII and junctional tension are higher in NCCs than OCCs

While the depletion of MyoII from AJs and AJ corrugations occurred in all follicle cells between stages 6 and 9, we observed pronounced differences in levels of AJ and actomyosin components between different follicle cell populations. Specifically, between stages 6 and 9, NCCs retained high junctional levels of E-cad and β-cat, whereas levels strongly decreased in OCCs (Fig 3A,C,C’, Fig S3A). In contrast, AJ-associated proteins such as N-cadherin (N-cad) or Echinoid (Ed) did not specifically decrease in cells contacting the oocyte (Fig S3A). Further, MRLC and its active form (MRLC-1P) were specifically enriched in the apical-medial domain of cells positioned over nurse cells between stages 6 and 9 but, like E-cad and β-cat, levels dropped in OCCs (Fig 3B,D,D’, Fig S3B). Follicle cell specific expression of a fluorophore-tagged MRLC verified that the apical domain of follicle cells alone reflected the differences in fluorescence intensity (Fig 3E-F’, Fig S3C). In contrast, levels of basal MyoII were significantly lower than apical levels (Fig 3F,F’’) and spatial patterns of basal MyoII did not correlate with nurse cell or oocyte contact (Fig S3D). Of note, anterior squamous-fated cells which are also in contact with nurse cells displayed a reduction in levels of AJ and MyoII components starting at stage 7/8 (Fig 3A-E). This correlated with apical expansion as a result of flattening, indicating dilution of apical and junctional proteins over a larger surface as cause for reduced protein levels. However, in contrast to anterior cells, decreasing levels of AJ and MyoII components in OCCs cannot be explained by dilution given the relatively smaller apical OCC surface area, if compared to NCCs. Combined, this suggests that nurse cell contact promotes maintenance of E-cad, β-cat and MyoII at the apical-junctional domain of NCCs. In agreement with this conclusion, high levels of apical-junctional E-cad/β-cat and MyoII also correlated with nurse cell contact in egg chambers containing a mislocalized oocyte (Fig 3G-G’’, Fig S3E-F’), confirming that oocyte and nurse cell contact rather than posterior terminal and main body patterning *per se* are sufficient to regulate apical-junctional E-cad/β-cat and MyoII levels.

**Fig 3:**
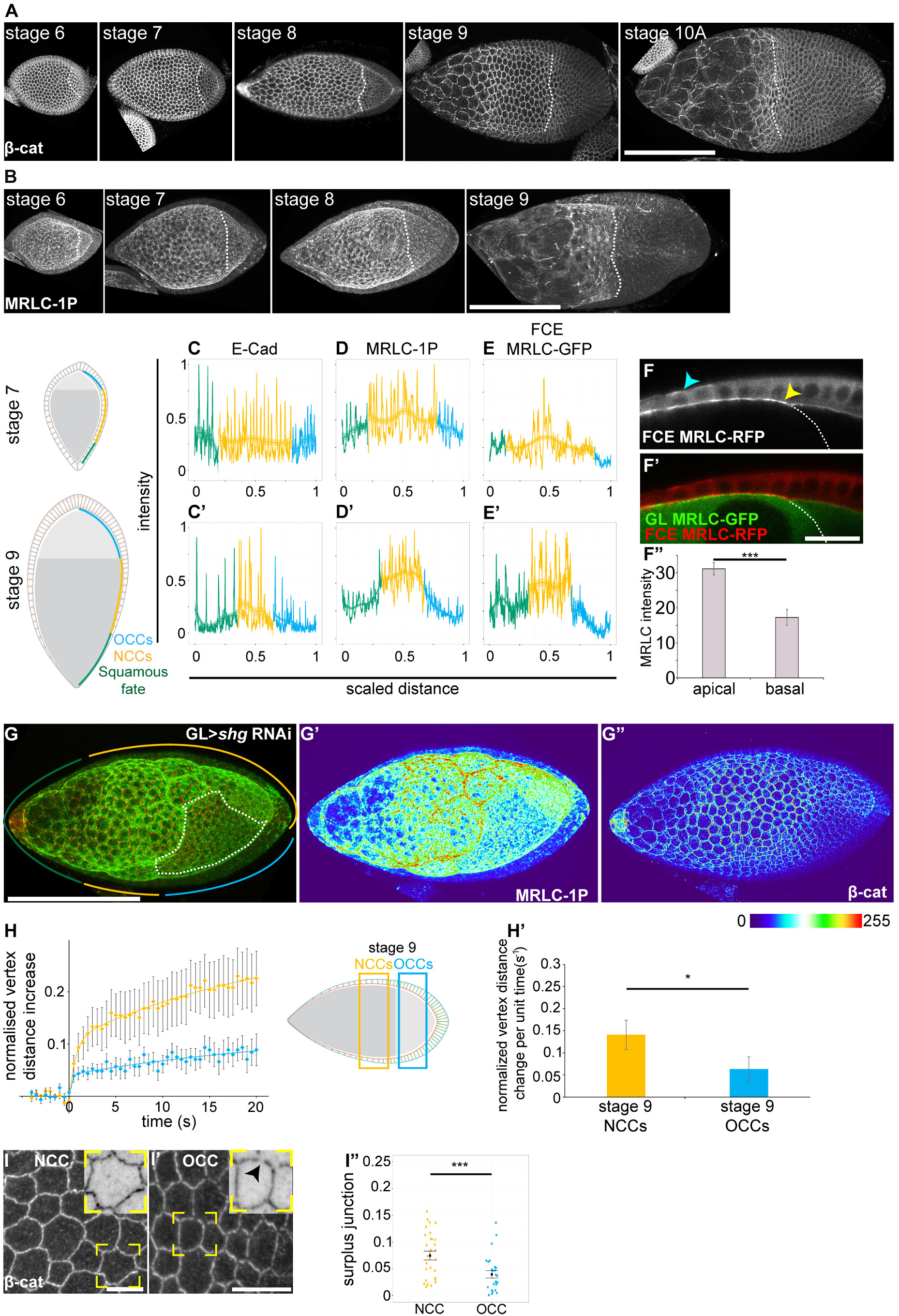
Levels of E-cad/β-cat, MyoII and junctional tension are higher in NCCs than OCCs. **(A,B)** Maximum projection of confocal sections to obtain *en face* view of AJs in egg chambers stained for β-cat and MRLC-1P. **(C-E’)** Representative line profiles of fluorescence intensities at apical-junctional domains in medial sections at stages 7 and 9. Intensities normalized to the maximum measured value were plotted for squamous cells (green), NCCs (orange) and OCCs (blue) along the length of an egg chamber scaled from 0-1 (anterior-posterior). Egg chambers were stained for E-cad (C,C’), MRLC-1P (D,D’) and FCE expression of *MRLC-GFP* (E,E’). For each marker, reproducible line profiles were obtained for n≥5 egg chambers. A fitted curve (see Exp. Proc.) is plotted along with standard error bounds, which aids in seeing intensity trends in different FCE populations. Note that peak intensities for E-cad coincide with junctions and decrease in OCCs. Peak intensities of MRLC coincide with junctional and medial positions. **(F-F”)** Medial section of an egg chamber with FCE expression of *MRLC-RFP* (F, red in F’) and germline expression of *MRLC-GFP* (green in F’). Yellow arrowhead points to the sharp drop in apical MRLC-RFP levels in the FCE at the oocyte boundary. Cyan arrowhead indicates basal MRLC. Graph displays mean±SEM of apical and basal MRLC fluorescence intensity in NCCs of egg chambers expressing *MRLC-GFP* or *MRLC-RFP* in the FCE only (F”). n=5 egg chambers. A t-test was performed. *** indicates p-value≤0.001. **(G-G”)** Maximum projections of confocal sections to visualize AJs (β-cat, red in G, thermal LUT in G’’) and MRLC-1P (green in G, thermal LUT in G’) in a stage 9 egg chamber. The germline (GL) expresses RNAi targeting *E-cad (shotgun, shg)* resulting in a mispositioned oocyte (framed by white dotted lines). Orange (NCC), blue (OCC) and green (squamous-fated) lines indicate different FCE populations. **(H,H’)** Graph (H) displays the normalized average increase in distance between vertices upon laser ablation of AJs (t=0) as a function of time in stage 9 NCCs (orange) or stage 9 OCCs (blue). Graph (H’) shows the normalized initial vertex distance change per unit time (see Exp. Proc.). Graphs display mean±SEM. n=9 each, two-sample t-test was performed. * indicates p-value≤0.05. Egg chamber scheme illustrates position of follicle cells subject to laser cuts. **(I-I”)** *En face* junctional sections at stage 9 in NCC (I) and OCC (I’) stained for β-cat. Insets show higher magnifications of yellow squares to visualize discontinuities and corrugations in AJs (arrowhead). Quantification of surplus junction length in NCCs and OCCs from one representative egg chamber (n=29 NCCs, n=26 OCCs) (I”). Graphs display mean±SEM. Statistically significant differences were obtained for 3 additional egg chambers (not shown). A WMW-test was performed. *** indicates p-value≤0.001. Scale bars (A,B, G-G”) =100μm, (F-F’)=20,(I-I”)=10 μm

To understand if differences in junctional E-cad/β-cat and medial MyoII levels between NCCs and OCCs correlated with differences in junctional tension, we analyzed vertex recoil velocities upon laser ablation in NCCs and OCCs at stage 9. Indeed, junctional tension in NCCs was higher than in OCCs (Fig 3H-H’). Importantly, junctional tension in NCCs was dependent on MyoII contractility. A RNAi mediated knockdown of the MRLC kinase *Rok* in the epithelium caused a significant decrease in the measured initial recoil velocities after junctional ablation at stage 9 (Fig S2G-G’’). Thus, high levels of medial MyoII in NCCs likely translate into higher levels of AJ tension if compared to OCCs. If higher levels of medial MyoII are the source of higher AJ tension in NCCs, then NCC AJs may be more corrugated than OCC AJs, as NCC AJs deflect more strongly by punctuate links to a contractile medial cortex. Indeed, we found that the surplus junctional length between NCC vertices is longer than between OCC vertices. Our results demonstrate that while overall AJ tension decreases between stage 6 and 9, NCCs maintain higher levels of AJ tension than OCC. While the overall decrease in AJ tension correlated with the medial shift of MyoII between stage 6 and 9, the relatively higher AJ tension in NCCs correlated with the presence of higher levels of medial MyoII and more pronounced AJ corrugations, if compared to OCCs. Combined, these observations support a model where medial MyoII controls tension at NCC and OCC AJs.

### Regulators of actomyosin contractility are required to prevent excessive NCC flattening

Our observations that high levels of medial MyoII, AJ tension and corrugations correlated with nurse cell contact at stage 9 contradicted expectations about the regulation of cell shape transition between cuboidal NCCs and columnar OCCs. Intuitively, a reduction in apical surface area of columnarizing OCCs in response to contact with the growing oocyte is expected to depend on relatively higher levels of apical-junctional contractility, whereas relatively larger apical surface areas in cuboidal NCCs in contact with nurse cells could be associated with reduced apical-junctional contractility.

Thus, to understand the functional relevance of higher levels of medial MyoII and junctional tension in NCCs, we genetically manipulated regulators of actomyosin contractility. Using RNAi-mediated knock-down driven by TJ-GAL4, a driver displaying highest activity after stage 5, we reduced *Rok* or *sqh* function in the entire epithelium. Strikingly, while OCCs displayed only minor alterations to cell shape at stage 9, mutant NCCs responded with an expansion of apical surface area and a reduction in lateral heights (Fig 4A-H’, Fig S4A-C’). Apical expansion of mutant NCCs was not due to loss of cells or multinucleation observed when cells lose MyoII function in mitotic stages 2 to 5 [20]. In fact, at stage 9, *Rok* RNAi expressing epithelia only rarely contained binucleate cells and the total number of cells was conserved (31.0±0.3 and 30.6±0.2. cells in a row along the A/P egg chamber axis in wild type and *Rok* RNAi expressing egg chambers, respectively, n=5 each). Importantly, the observed increase in apical areas of NCCs lacking *Rok* or *sqh* function occurred at the expense of corrugation (Fig 4C,D, Fig S4D,D’) supporting the idea that corrugations are maintained by medial MyoII activity tethered to point contacts at AJs. Combined, these results demonstrate a specific requirement for higher levels of medial MyoII and AJ tension in maintaining NCC shape by constraining apical NCC areas and thus preventing flattening. Strikingly, OCCs did not exhibit a similarly strong requirement for MyoII activity demonstrating that maintenance of apical OCC areas does not depend on high levels of actomyosin contractility.

**Fig 4:**
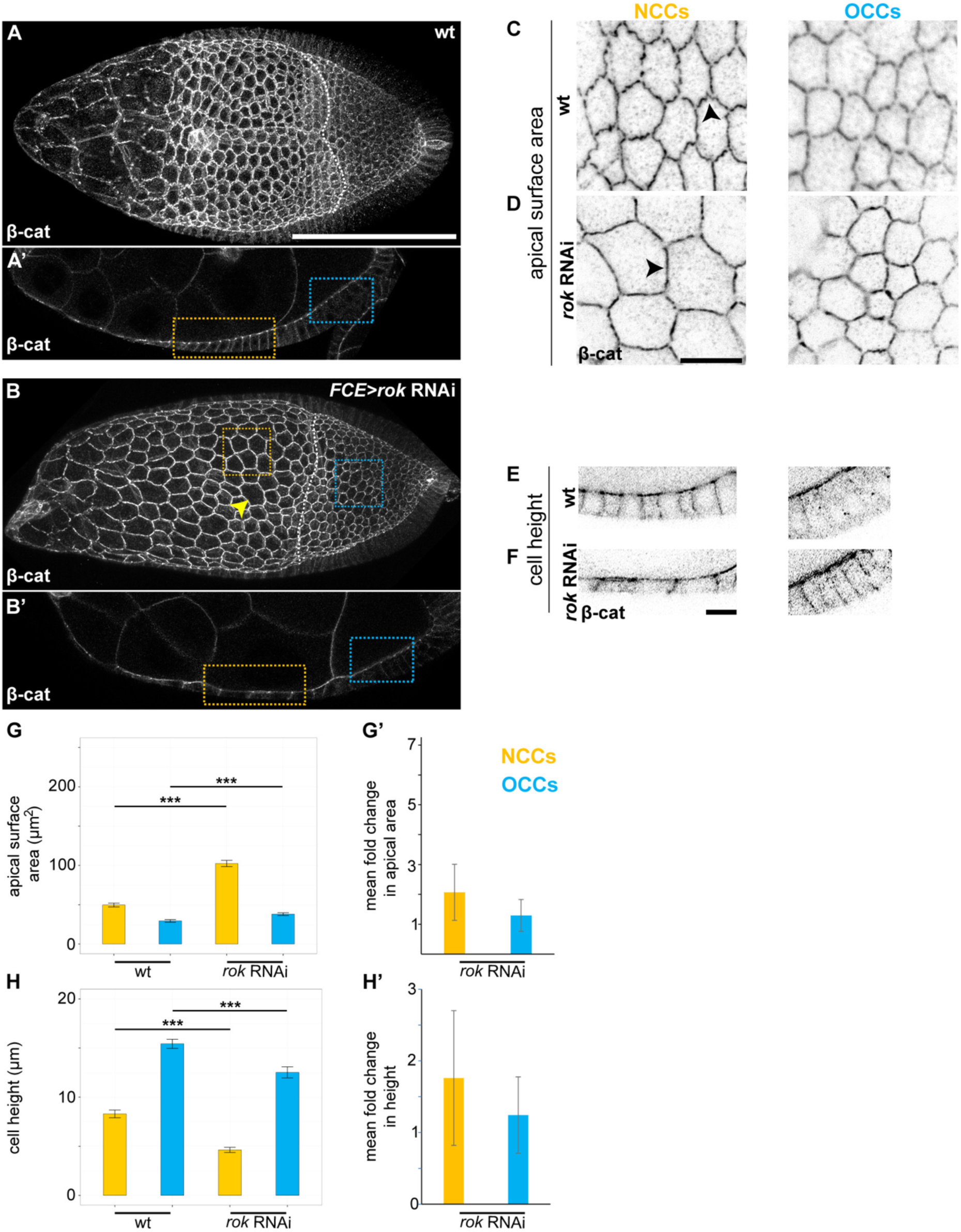
Regulators of actomyosin contractility are required to prevent excessive NCC flattening. **(A-F)** Maximum projections for *en face* view of AJs (A,B,C,D) and medial sections (A’,B’,E,F) of wild type (wt) egg chambers or one with FCE expression of *rok* RNAi stained for β-cat. Yellow (NCC) and blue (OCC) dotted lines (A’-B’) frame apical cell areas (C,D) and cell heights 1^(E,F). Black arrowheads indicate corrugations in wt (C) and reduced corrugations in *rok* RNAi (D) NCCs. Yellow arrowhead in (B) points to a multinucleate cell. Such cells were rare and were excluded from the quantification of cell areas and cell numbers. **(G-H’)** Mean apical areas (G) and cell heights (H) ±SEM and the mean fold change relative to wt in areas (G’) or heights (H’) with standard errors computed by propagation of error for NCCs (yellow) and OCCs (blue) upon FCE expression of *rok* RNAi. See Table S2 for sample sizes. Welch t-tests were performed. *** indicates p-value≤0.001. Scale bar (A-B’)=100μm, (C-F)=10μm

### Regulators of AJ length are required to prevent excessive NCC flattening

To understand if AJ function also controls the size of apical NCC surfaces between stages 6 and 9, we genetically manipulated core components of AJs. Mosaic analysis of *E-cad* null clones revealed minimal changes to cell shape, because of compensatory upregulation of N-cad (data not shown) [38]. We thus expressed *α-catenin (α-cat)* RNAi in the entire epithelium or generated *β-cat* mutant mosaic clones. Both approaches eliminate the catenin-mediated linkage of E-cad or N-cad to Actin and thus AJ formation [5, 6, 39]. Using this strategy, we found that specifically *α-cat* and *β-cat* mutant NCCs but not OCCs flattened with domed apical membranes by stage 9 (Fig 5A-B’,D,E,G,H, Fig S5A-D), demonstrating that AJs are principally important to mediate the function of high medial MyoII and AJ tension in NCCs.

**Fig 5:**
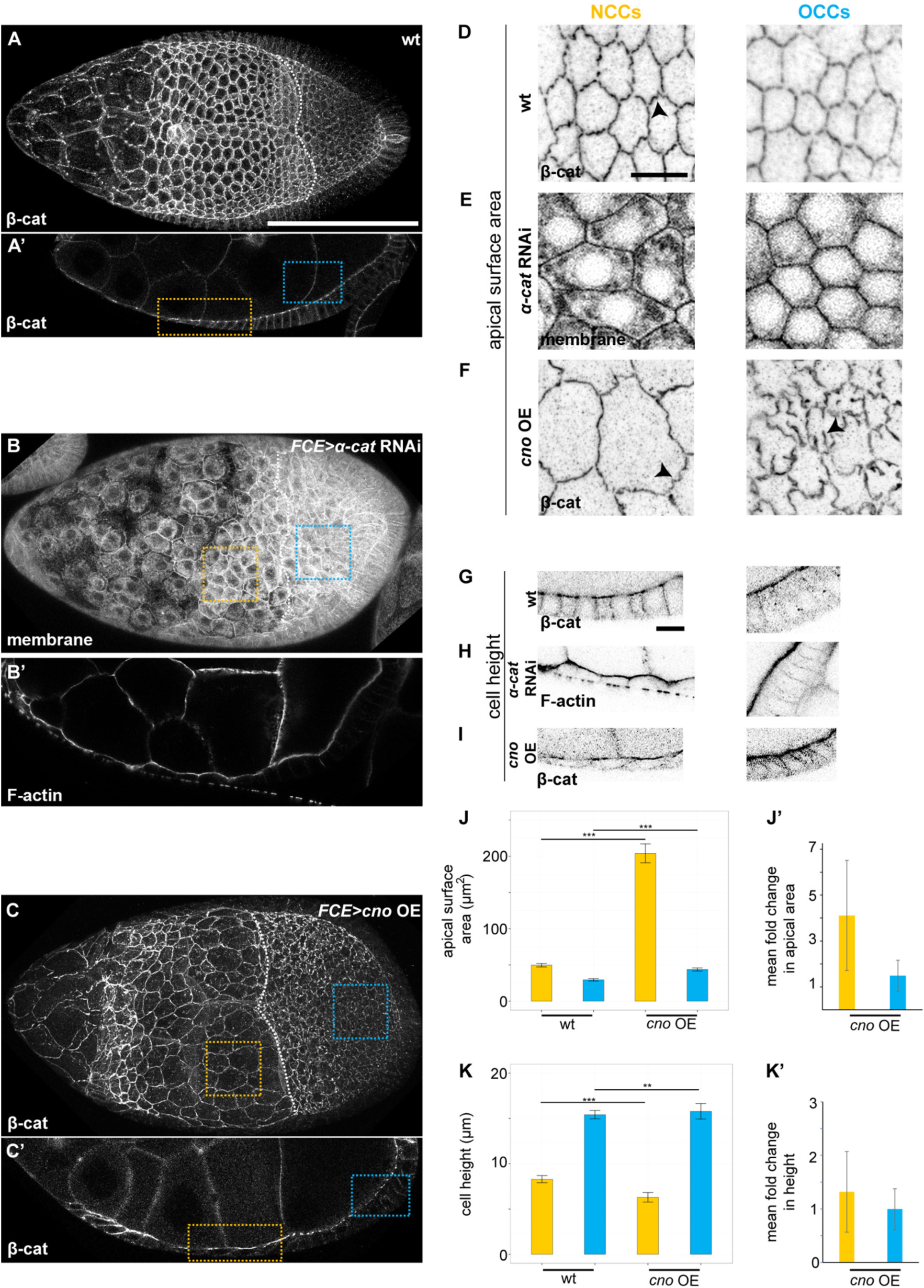
Regulators of AJ length are required to prevent excessive NCC flattening. **(A-I)** Maximum projections for *en face* view of AJs (A,B,C,D-F) and medial sections (A’,B’,C’,G-I) of wt egg chambers or those with FCE expression of *α-cat* RNAi or overexpression of *cno* (OE) stained for β-cat. Yellow (NCC) and blue (OCC) dotted lines (A’-C’) frame apical cell areas (D-F) and cell heights (G-I). Black arrowheads (E,F) indicate corrugations in wt and hypercorrugations in *cno* OE cells. **(J-K’)** Mean apical areas (J), cell heights (K) ±SEM and mean fold change relative to wt in areas (J’) or heights (K’) with standard errors computed by propagation of error for NCCs (yellow) and OCCs (blue) upon *cno* overexpression in the epithelium. See Table S2 for sample sizes. Welch t-tests were performed. *** indicates p-value≤0.001. Scale bar (A-C’)=100μm, (D-I)=10μm

As the elimination of catenin function prevents cell-cell adhesion and therefore the transmission of forces within the junctional network, we wanted to analyze known regulators that modulate rather than define AJ function. Strikingly, we found that targeted overexpression of the AJ regulator Afadin (Canoe (Cno)) [40-42] caused dramatic flattening of NCCs by stage 9 (Fig 5C,C’,F,I-K’) despite the presence of apical-medial MyoII in *cno*-expressing NCCs (Fig S5E,F). In contrast to NCCs, *cno*-expressing OCCs maintained their relatively small apical areas and columnar shape but displayed severe AJ hypercorrugation. Thus, Cno overexpression lead to an increase in AJ length and suggested that this surplus AJ length caused apical NCC expansion and cell flattening. Overexpression of a GFP-tagged Cno revealed that Cno localized with E-cad in vesicle-like structures at AJs (Fig S5G), suggesting that it modulates E-cad trafficking. Importantly, apical areas of *cno*-expressing NCCs were larger than those observed in *sqh* RNAi or *Rok* RNAi expressing NCCs (Fig 4G, 5J). This was not due to reduced follicle cell numbers enveloping the germline surface area (31.0±0.3 and 30.4±0.4 cells in a row along the A/P egg chamber axis in wild type and *cno*-expressing egg chambers, respectively, n=5 each). We thus suggest that absolute AJ length defines the maximum size of an apical NCC area. This maximum possible NCC area is reduced by medial MyoII to a smaller corrugated surface, whose size depends on a functional ratio between medial contractility and AJ length.

In our search for additional modulators of apical NCC area, and thus cuboidal shape, we found that targeted overexpression a dominant-negative *Rac1 (Rac1^DN^)* [43] caused NCCs to flatten more extensively than OCCs (Fig S5H-I’). Apical expansion of *Rac1*^*DN*^ expressing NCCs was associated with abnormal tubular AJs protruding into the apical surface, indicating that Rac1 also modulates corrugated AJ architecture (Fig S5I). Moreover, in agreement with coordination of AJ function by apical polarity determinants [1], we found that reducing levels of the apical polarity proteins aPKC or Crumbs (Crb) in the epithelium caused pronounced NCC flattening (Fig S5J-M). Combined these results demonstrate that the maintenance of apical NCC areas, and thereby of cuboidal shape, relies on the precise regulation of AJ organization and, importantly, length. In contrast, apical OCC areas and thereby columnar shape did not exhibit a similarly strong requirement for AJ function, suggesting that NCCs and OCCs exhibit very different molecular requirements for the maintenance of their apical areas and associated cell shapes.

### Maintenance of apical NCC surface area through regulation of AJ length and contractility is required to complete cuboidal-columnar shape transitions

To assess the organ-level consequences of the specific sensitivity of NCCs to deregulation of MyoII and AJ function, we closely analyzed egg chambers with *sqh* RNAi, *Rok* RNAi, *cno* or *Rac1*^*DN*^ expressing epithelia. Whereas *sqh* RNAi and *Rok* RNAi expressing chambers progressed to late stages of development, all *cno* and *Rac1*^*DN*^ expressing egg chambers degenerated by what at first glance appeared to be stage 9. Specifically, anterior cells had flattened and main body NCCs were still positioned over nurse cells, indicative of developmental stage 9. However, we found that chamber sizes were unusually large (Fig S6). To eliminate that this was just a coincidence in degenerating egg chamber, we analyzed the size of viable *cno* and also of *Rok* RNAi expressing egg chambers with stage 9 morphologies. Importantly, the size of their wild type germline was significantly larger than of stage 9 wild type chambers, and more similar to a size normally observed at stage 10A (Fig 6A-D). To test if the germline continues to grow by maintaining a nurse cell/oocyte ratio characteristic of stage 9, or if this ratio also advanced to stage 10A, we measured nurse cell and oocyte sizes. Indeed, the nurse cell/oocyte ratio in egg chambers with stage 9 *Rok* RNAi or *cno* expressing epithelia was closer to that of stage 10A wild type egg chambers, demonstrating that oocytes expanded normally with germline size. Combined, this demonstrates that the development of the germline continues normally. However, flattening of *cno* and *Rok* RNAi expressing NCCs expands the total NCC surface anteriorly and beyond the reach of the normally expanding oocyte (Fig 6F-F’’). The severe expansion of *cno*-expressing NCCs ultimately causes a failure of all main body cells to ever acquire oocyte contact, likely underlying egg chamber degeneration. In conclusion, maintenance of apical NCC areas is critical to facilitate contact with the expanding oocyte, and thus to complete cuboidal to columnar cell shape transitions at stage 9.

**Fig 6:**
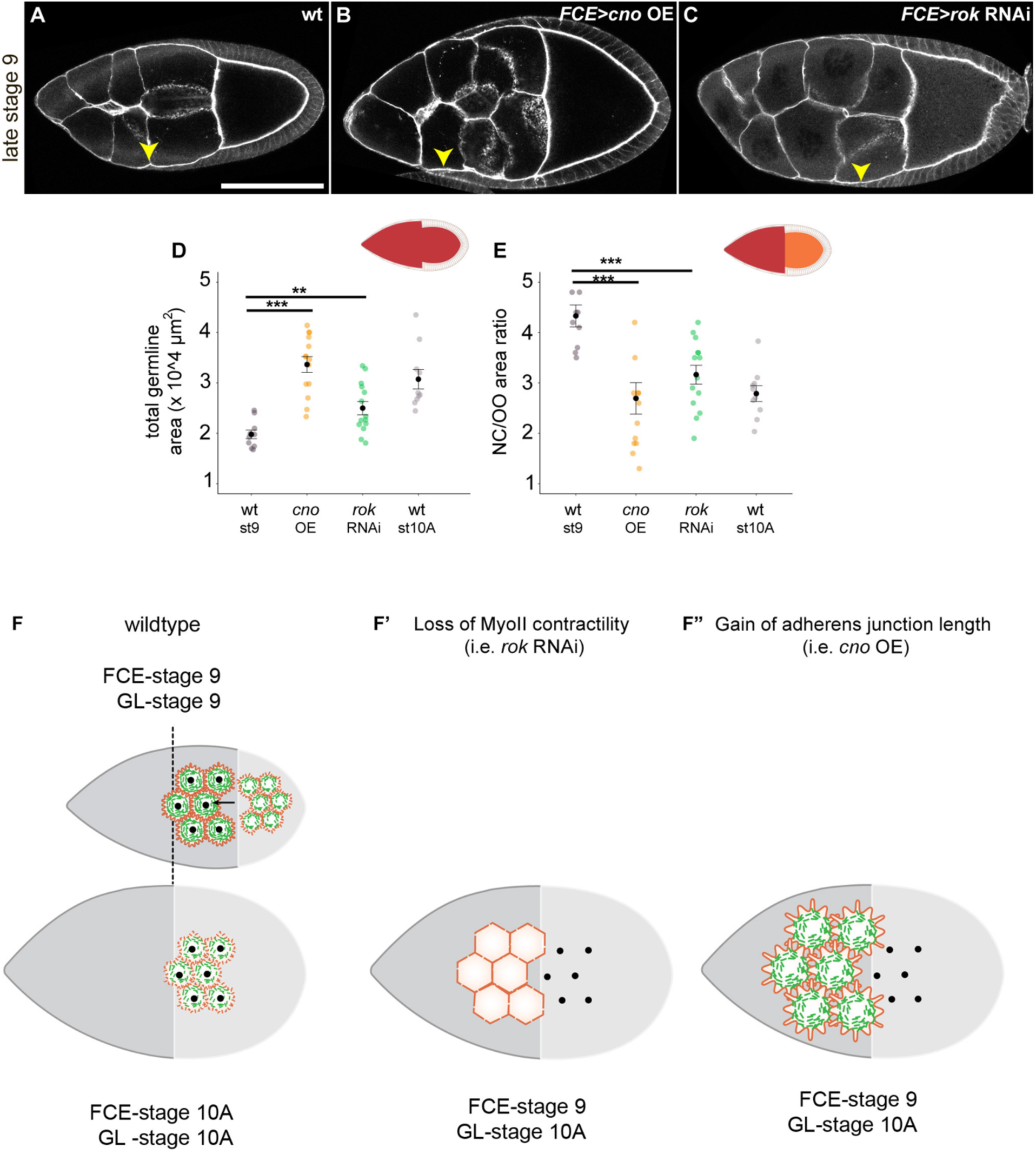
Maintenance of NCC cuboidal shape through regulation of AJ length and contractility is required to complete cuboidal-columnar shape transitions. **(A-C)** Medial sections of morphological late stage 9 wt egg chambers (indicated by presence of FCE cell shape gradient) (A), or with FCE expression of *cno* (B) or *rok* RNAi (C) stained for F-actin and β-cat. Yellow arrowheads point to most anterior NCCs. **(D,E)** Total germline area (D, dark red in scheme) or nurse cell-to-oocyte area ratios (E, ratio of dark red to orange area in scheme) were calculated in medial sections of wt egg chambers (stage 9 or 10A) or egg chambers with FCE expression of *cno* or *rok* RNAi. n≥8 egg chambers for each genotype. Welch t-tests were performed. *** indicates p-value≤0.001, ** indicates p-value≤0.01 **(F-F”)** In a wild type egg chamber, the oocyte grows anteriorly (black arrow) to acquire its stage10A size (dotted black line shows where the oocyte boundary will be at stage 10A). Thus, NCCs at stage 9 become OCCs at stage 10A (see also Fig 1E). Corrugating FCE junctions are labelled in orange and medial MyoII in green. Black dots label the central position of apical NCCs surfaces at stage 9 and 10A. Note that their absolute position does not change but the oocyte expands underneath them. Upon loss of MyoII function in the FCE, NCCs expand and lose AJ corrugations (F’). Upon gain of AJ length in the FCE, excess junction length promotes apical NCCs expansion (F”). Apical expansion of mutant NCCs in (F’, and F’’) displaces their centre positions anteriorly. The oocyte and the total germline (GL) grow to a stage 10A size but the FCE remains in contact with nurse cells, characteristic of stage 9. Black dots in F’,F” label where NCCs should have been had they maintained normal apical surface size. Scale bar (A-C)=100μm

### Apical-junctional NCC contractility promotes nurse cell cluster elongation

Compared to OCCs, NCCs responded more sensitively to the manipulation of actomyosin and AJ function by expanding their apical surface areas by stage 9. This implies that contact with nurse cells drives apical surface area expansion in NCCs. Nurse cells may drive expansion by coordinating surface growth or surface shape with overlying NCCs. A 1.8-fold increase in nurse cell surface contributes to germline growth between stage 8 and 10A when cell shape transitions primarily occur (Kolahi et al, 2009). Thus, the up to 5-fold increase in apical areas of *cno*-expressing NCCs is not sufficiently accounted for by growth alone.

We thus asked whether nurse cell shape may also drive apical expansion of mutant NCCs. Strikingly, apical expansion in *cno* or *Rok* RNAi expressing epithelia coincided with bulging of individual nurse cells into the apical surface of NCCs at stage 9 (Fig 7A,B). The deformation of the apical surface furthermore coincided with a significant widening at the dorsal-ventral (D/V) axis of the nurse cell cluster, concomitant with a shortening of the A/P axis (Fig 7C). Significant differences in the aspect ratio of total egg chambers containing *cno* or *Rok* RNAi expressing epithelia was not observed at stages 7 and 8 (Fig S7A). This excludes defects acquired during rotation-driven axis elongation of egg chambers up to stage 8 as cause of nurse cell cluster aspect ratio changes at stage 9. Combined, this suggested that apical NCC surfaces constrain bulging of individual nurse cells and widening of the D/V axis of the entire nurse cell cluster at stage 9.

**Fig 7:**
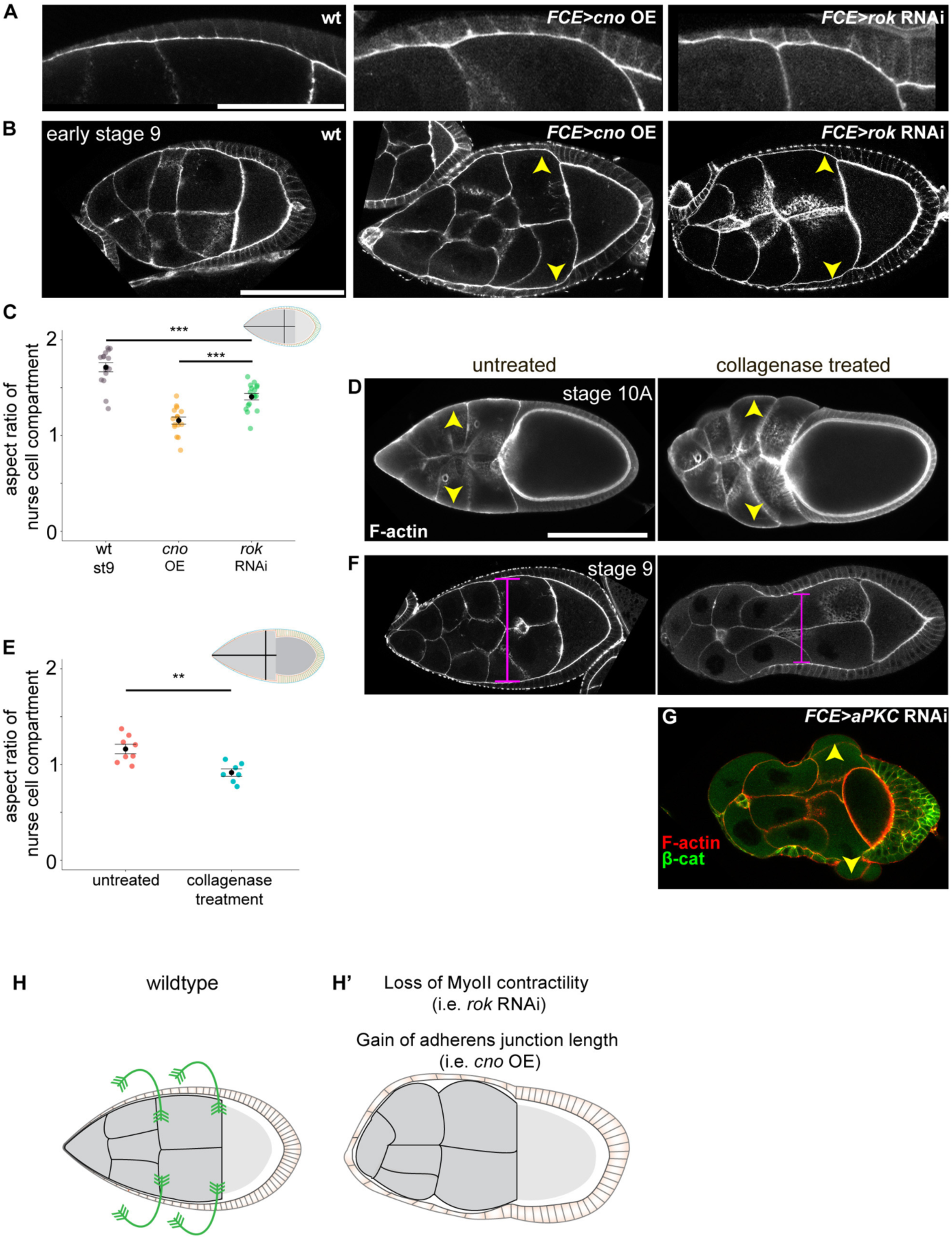
Apical-junctional NCC contractility promotes nurse cell cluster elongation. **(A)** Bulging of individual nurse cells upon FCE expression of *cno* or *rok* RNAi compared to a wt egg chamber stained for F-actin and β-cat. **(B)** Medial sections of stage 9 egg chambers, either wt or with FCE expression of *cno* or *rok* RNAi stained for F-actin and β-cat. Yellow arrowheads point to nurse cell cluster widening at NCC positions. **(C)** Length-to-width aspect ratio of nurse cell compartments (see scheme) for wt egg chambers or with FCE expression of *cno* or *rok* RNAi at stage 9. n≥9 egg chambers for each genotype. A t-test was performed. *** indicate p-value≤0.001. **(D,F,G)** Medial sections of egg chambers untreated or treated with collagenase at stage 10A or stage 9 (F,G) stained for F-actin and β-cat. Egg chambers express *vkg-GFP (Viking, vkg,* CollagenIV) (D) or *aPKC* RNAi in the epithelium (G). Yellow arrowheads point to nurse cell bulging upon collagenase treatment (D,G). Magenta lines in F indicate the DV axis width of the nurse cell compartments. **(E)** Length-to-width aspect ratio of nurse cell compartments of untreated (n=8) and collagenase treated (n=7) stage 10A egg chambers. A t-test was performed. ** indicate p-value≤0.01. Scale bars (A)=50 μm, (B-G)=100μm **(H)** Loss of MyoII function or gain of AJ length (H’) reduces circumferential constraints imposed by the apical NCC surface on nurse cell cluster rounding (H).

To provide additional evidence for this idea, we investigated what shape nurse cells acquired in the complete absence of external constraints. To this end, we enzymatically removed the basement membrane, thought to contribute to elongated egg chamber shape [24], from stage 10A egg chambers with collagenase. At this stage, nurse cells are only covered by ultrathin squamous cells, which are not expected to contribute significantly to the combined material properties of the nurse cell-epithelial cell interface. Indeed, we found that removal of the basement membrane caused bulging of individual nurse cells and reduction of the nurse cell cluster aspect ratio to that of a much round shape (Fig 7D,E, Fig S7B). This demonstrates that, in the absence of external constraints, the default shape of the nurse cell cluster is round rather than elongated.

To test if NCCs compressed the D/V axis of the nurse cell cluster at earlier stages, we enzymatically removed the basement membrane from stage 9 egg chambers. Here, we observed nurse cell bulging only at the anterior pole where cells are flat. More importantly, however, this coincided with circumferential constriction of egg chambers at NCC positions (Fig 7F). This indicates that NCCs exert circumferential contractility on the nurse cell cluster, which otherwise would have expanded in D/V into a rounder shape if unconstrained (compare to Fig 7D). As a consequence of NCCs circumferentially squeezing nurse cells, nurse cells bulge where external constraints like the basement membrane are removed. To provide additional evidence for the idea that the nurse cell cluster’s D/V axis is constrained by NCCs contractility, we eliminated NCCs and assessed unconstrained nurse cell shape also at stage 9. RNAi-mediated knockdown of the apical determinant aPKC caused extreme NCC flattening. Upon additional removal of the basement membrane, we observed pronounced bulging of nurse cells, which flat mutant NCCs failed to constrain (Fig 7G). Thus, NCCs actively constrain nurse cell and nurse cell cluster shape at stage 9. Accordingly, nurse cell shape did not change at all when collagen encapsulating stage 8 egg chambers was removed (Fig S7C), demonstrating that when anterior cells have not flattened yet, the follicle epithelium is sufficient to constrain nurse cell cluster shape. Combined, our data demonstrates that reinforced contractility and regulation of AJ length in the apical surface of NCCs suppresses rounding of individual nurse cells and imposes circumferential constriction on the nurse cell cluster to ensure elongation during oocyte growth at stage 9 (Fig 7H,H’).

## Discussion

### Regulation of tensile stress in the medial-junctional network

In this study, we investigate how tensile stress within the AJ network of a closed epithelial sheet integrates growth of a neighbouring tissue, mediates cell shape transitions and channels growth into organ elongation. Surprisingly, overall AJ tension decreases between stages 6 to 9, despite the expectation that growth of the germline surface stretches and thus increases tension in the overlying epithelium. Among other possibilities, for example changing expression of MyoII regulators [44, 45], the observed decrease in AJ tension may arise by a shift to medial contractility acting at an angle to AJs. Medial contractility acting at an angle to AJs is expected to reduce the effective force felt by cellular vertices if compared to the same amount of contractility acting in parallel to AJs. We do not know which signal initiates elaboration of a medial actomyosin web or the onset of corrugations in AJs. Mechanosensing of germline growth may provide external cues for medial MyoII and AJ remodelling [46, 47]. However, we speculate that as AJ tension reduces, external forces can more easily deform the junctional network to assist total surface area expansion of the epithelium during growth of the germline. In support of this idea, squamous cell flattening (stage 6-10A) has been suggested to be mediated by apical relaxation promoting compliance to germline growth [11].

As has been described for medial MyoII oscillations driving ratchet-like apical constriction [5, 48, 49], we suggest that AJ corrugations arise by deflection of AJs into the medial plane due to high radial tension at AJs. Indeed, upon loss of *Rok* or *sqh* function, AJ corrugations disappear as the apical surface expands, demonstrating that medial MyoII constrains apical areas and maintains corrugations by linking to AJs. Moreover, stronger corrugations in NCCs than OCCs correlate with higher junctional tension, suggesting that corrugations are not just a consequence of surplus AJ length surrounding a limited apical surface but that corrugations arise by active tension imposed on AJs. However, hypercorrugated junctions in *cno*-expressing OCCs demonstrate that a surplus of junctional material can also promote corrugations. Importantly, the ratio of absolute junction length to medial contractility appears to regulate the size of the apical surface. Excessive apical expansion of *cno*-expressing NCCs occurs even as a contractile machinery is in place to counteract it. In contrast, the relatively smaller surface area expansion upon *Rok* or *sqh* appears limited by regular junctional length. This interpretation also explains the relatively milder tissue-level phenotypes observed upon *Rok* or *sqh* LOF, in comparison to *cno*-overexpression.

### Modulation of cuboidal cell shape by nurse cell contact

Despite the overall reduction in AJ tension between stages 6 and 9, we demonstrate that NCCs locally reinforce AJ contractility at stage 9. Moreover, NCCs respond more sensitively to the manipulation of actomyosin and AJ function than OCCs, even though main body NCCs and main body OCCs, specifically, are of the same fate. We thus suggest that MyoII enrichment in main body NCCs is a local response to resist apical expansion driven by nurse cell rounding and growth. This ensures that NCCs conserve their apical surface size, and consequently maintain cuboidal shape and relative position to allow contact with an expanding oocyte. We expected that the OCCs would display higher junctional tension than NCCs to create the relatively smaller apical areas characteristic of columnar OCCs or a higher apical stiffness predicted by [11]. Instead, we found that OCCs display reduced levels of MyoII, junctional tension and lower sensitivity to the loss of MyoII and AJ function, if compared to NCCs. On one hand this indicates that the aspect ratio change during columnarisation is not driven by an increase in OCC AJ contractility relative to NCCs. Thus, columnar shapes do not solely arise as a consequence of intrinsic contractility at apical or basal cell domains [f.e. 50]. Instead, we speculate that OCCs may be subject to weaker external forces arising from oocyte growth and thus acquire the small apical areas associated with columnar shape. Over nurse cells, this fate-specified columnar shape is stretched into a cuboidal aspect ratio, depending on differential interactions of the apical epithelial surface with nurse cells and the oocyte. We speculated that direct adhesion between NCCs and nurse cells could drive apical expansion during coordinated growth. However, RNAi mediated double-knockdown of N-cad and E-cad in the germline inhibited the migration of border cells, as previously reported [51], but did not disrupt epithelial shape transitions (data not shown). Therefore, Cadherin-dependent adhesion between NCC and nurse cells cannot account for NCC-specific behaviors and future studies need to address other mechanisms of NCC-nurse cell communication.

### Organ elongation by circumferential apical contractility

Previous studies suggest that elongated egg chamber shape is determined by a molecular corset at the basal epithelial surface channelling growth of the egg chamber into the A/P axis [14, 24, 25, 29, 52]. Polarized ECM properties act between stage 2 and 7 [14, 24, 25] and basal actomyosin contractions ensure egg elongation from mid 9 to 10B [29]. Our study suggests a mechanism that ensures nurse cell cluster and thus egg chamber elongation during stages 8 to 9. Our data is consistent with a model where relatively higher levels of apical-junctional NCC contractility establishes a radially contractile sleeve constraining nurse cell bulging and nurse cell cluster rounding in the D/V egg chamber axis. Accordingly, genetic reduction of apical-junctional contractility or an increase in AJ length causes nurse cell bulging and nurse cell cluster rounding while NCCs expand and flatten. Importantly, patterned apical contractility also shapes the egg chamber prior to stage 6 [30]. Thus, the critical importance of the apical domain prior to stage 6 and at stages 9 suggests that basal and apical constraints imposed on egg chamber shape provide alternating mechanisms for egg chamber elongation at different stages of egg chamber development.

## Experimental Procedures

### *Drosophila* stocks and genetics

All experiments were performed on *Drosophila melanogaster*. For detailed genotypes listed for each figure, please refer to Table S1. Stocks and experimental crosses were maintained on standard fly food at 18°C or 25°C. Mosaic analysis was performed using the FLP/FRT and the actin-flip-out system [53]. For follicle epithelium clones, FLP expression was induced in young adult females using a heat shock for 1 h at 37°C. For germline clones, FLP expression was induced for 1 h at 37°C at 96 h and 120 h after egg lay at 25°C. Flies were fed yeast paste for 48 to 72 h before dissection.

### Immunohistochemistry and imaging

Ovaries were dissected and fixed in 4% formaldehyde/PBS for 15 min at 22°C. Washes were performed in PBS + 0.1% Triton X-100 (PBT). Ovaries were incubated with primary antibodies in PBT overnight at 4°C: guinea-pig anti-Spaghetti-squash 1P (MRLC-1P) (1:400, gift from Robert Ward), mouse β-catenin (1:100, DSHB, N27A1), rat anti-E-cadherin (1:100, DSHB, DCAD2), rabbit anti-GFP (1:200, Thermo Fisher, G10362), rat anti-RFP (1:20, gift from H. Leonhardt, 5F8), Dlg (1:100, DSHB, 4F3), rat anti N-Cad (1:20, DSHB, DN-EXH8), mouse anti-PKC ζ (1:50, Santa Cruz, H-1,sc-17781), mouse β-gal (1:1000, Promega Z378B). Ovaries were incubated with secondary antibodies (coupled to Alexa Fluorophores, Molecular Probes) for 2 h at 22 °C. DAPI (0.25 ng/μl, Sigma), Phalloidin (Alexa Fluor 488 and Alexa Fluor 647, 1:100, Molecular Probes, or Phalloidin-TRITC, 1:400, Sigma). Egg chambers were mounted using Molecular Probes Antifade Reagents. Samples were imaged using Leica TCS SP5, SP8 or ZeissLSM880 confocal microscopes. Samples were processed in parallel and images were acquired using the same confocal settings, if fluorescence intensities had to be compared. Super-resolution imaging was performed using an Airyscan detector on a Zeiss LSM880 confocal microscope and images were post-processed with ZEN [54]. Images were processed and analyzed using FIJI (ImageJ 1.48b) [55].

### Live imaging

Individual ovarioles were dissected out of the muscle sheet and were mounted with a minimal volume of Schneider’s medium supplemented with FBS and insulin as described in [56] on a standard microscope slide with spacers fashioned from double-sided tape, covered with a coverslip and sealed with Halocarbon oil. Super-resolution imaging was performed using an Airyscan detector on a Zeiss LSM880 confocal microscope and post-processed with ZEN [54]. Images were acquired at a 30 s time interval.

### Collagenase treatment

Individual ovarioles were dissected from the surrounding muscle sheet and were incubated in Schneider’s medium supplemented with 1000 Units/ml collagenase (CLSPA; Worthington Biochemical Corp) for up to 30 min, rinsed in 1X PBS three times and then fixed and immunostained individually as described above in an 8-well tissue culture dish.

### Laser ablation

Laser ablation on live egg chambers expressing Shg-GFP [57] were performed on two set ups – using the inverted microscope set up described previously [36] (Fig 2 and 3) or an inverted Zeiss Spinning Disc (Yokogawa CSU-22) with a laser ablation unit (Rapp OptoElectronic) (Fig S3). Briefly, individual ovarioles were dissected out of the muscle sheet and were mounted on a standard microscope slide with spacers fashioned from double-sided tape, covered with a coverslip and sealed with Halocarbon oil (Sigma). Experiments were performed on freshly dissected ovarioles prepared every 20 minutes. 32 pulses/µm of the laser (λ=355nm) at 1000 Hz was applied at a length of 0.22 µm for ablations of cell-cell junctions (Fig 2 and 3). Images were taken every 0.3 s (Fig S3) or 0.5 s (Fig 2 and 3) for up to 40 s.

### Image Analysis and Quantification using FIJI

All images and movies were analyzed in FIJI (ImageJ 1.48b) [55], unless otherwise stated. Graphs were generated with Microsoft Excel 365 or R version 3.2.0. Statistical tests were performed in R 3.2.0. Data sets were checked for normality of distribution with Shapiro’s test and homogeneity of variances by applying Bartlett’s or Levene’s test. Statistical tests are indicated in figure legends. The α value for statistical analysis was set to 0.05 (α = 0.05).

#### Fluorescence intensity quantification

Measurement of fluorescence intensity traces for junction and cytoskeleton markers (Fig 2,3,S2,S3) were performed using line and profile plot tools in FIJI. The surface occupied by squamous fated cells was approximated by a line of the same length as which was obtained for OCCs in the same egg chamber. The remaining segment between ‘squamous-fated’ and OCC cells was denoted as NCCs. A fit was applied to the intensities using a smoothing function in R which automatically chooses a curve fitting method based on the group of the largest size of data points between squamous fated cells, NCCs or OCCs for each stage. Apical and basal MRLC intensity in the NCCs was measured in mid sections of egg chambers with the line tool in FIJI and subtracting the background intensity.

#### Quantification of apical cell areas, cell heights and AJ corrugations

Apical areas of epithelial cells were measured at the level of AJs using the polygon tool. Heights were measured using the line tool in a medial cross-section of the egg chamber. Junctional corrugations (surplus junction length) were quantified by forming a ratio of (1) junction length obtained by tracing the β-cat signal between two vertices using the segmented line tool and (2) the distance between the same vertices obtained by using the straight-line tool. This value would theoretically = 1 when the junction is a straight line and >1 when (1) > (2).

#### Analysis of vertex displacement and initial recoil velocities after laser ablation

To measure vertex displacement after ablation of AJs, a kymograph of the AJs between the two vertices was generated in FIJI. The vertices of the ablated junction were tracked pre-and post-ablation and distances between the vertices were obtained for each time point over the period of recording. For each ablation event, the change in distance between the vertices at any post-ablation time point relative to the average distance from 10 pre-ablation time points was obtained. The change in distance was normalized to the average junction length across all samples within one experimental condition. Finally, the mean relative distance was plotted as a function of time. In a first approach, a double exponential fit [36, 58] was applied to estimate the initial velocity of the average curves: d(t) = d1(1-e-t/T1) - d2(e-t/T2 - e-t/T1), where T1 is the slow relaxation time and T2 is the fast relaxation time of the vertices of ablated cell bonds. d1 is the final change of distance between vertices of ablated cell bonds at t → ∞ and d2 is the change in distance due to fast relaxation only. The fit parameters were calculated and the standard error was determined as shown in Table S3. The fit parameters d1 and T1 are poorly estimated for some data sets by: d(t) = d1(1-e-t/T1) - d2(e-t/T2 - e-t/T1) (Table S4). Fast time scale responses are associated with linear elastic behavior of the cytoskeleton cortex whereas slower ones with viscous behavior. Since, T2 or fast relaxation time ranges from 0.3 to 1 s in our measurements and is well estimated, we assume that the recoil of the vertices in this time interval to be like that of a linear elastic solid and thus the magnitude of initial velocity is directly proportional to the tension in the junctions. Thus instead of obtaining the initial velocity v0 by solving this equation: v0 = d1/T1 - d2 (1/T1- 1/T2) as described previously [58], we present initial velocities (Fig 2, 3 and S3) by calculating the slope of the curve between t=0 and t= 0.5 or 0.6 s which is expected to approximately cover the linear phase of the curves [59].

#### Aspect ratio measurements of nurse cell compartments and total egg chamber

Using the line tool in FIJI, the maximum width (W) across posterior nurse cells and the maximum length (L) of the nurse cell compartment or the total egg chamber measured from and to basal surfaces at the maximum width and length in a medial section was measured and the ratio of length to width was obtained.

#### Germline area and nurse cell-oocyte area ratio measurements

Using the Polygon tool in FIJI, the traces of the nurse cell compartment and oocyte were generated in the medial section of the egg chambers. Both areas were summed for total germline area and used as a proxy for volume of egg chamber. Ratio of nurse cell to oocyte for relative size was obtained.

## Author Contributions

Conceptualization RB, VW, AKC; Investigation RB, VW, MR, AKC; Writing RB, VW, AKC; Supervision AKC

## Acknowledgements

We thank G. Salbreux, S. Grill and the Life Imaging Center (LIC, University of Freiburg) for discussions and technical help with experiments. We thank R. Ward, Y. Bellaiche, E. Knust, S. Eaton, U. Tepass, M. Grammont, A. Carmena and H. Leonhardt for sharing reagents. We thank BDSC, VDRC and DSHB for providing fly stocks and antibodies. We thank the IMPRS-LS and SGBM graduate schools for supporting our students. Funding for this work was provided by the DFG (SPP1782).

